# SPAT: Surface Protein Annotation Tool

**DOI:** 10.1101/2023.07.07.547075

**Authors:** JF Spinella, L Theret, L Aubert, E Audemard, Geneviève Boucher, S Pfammatter, E Bonneil, ME Bordeleau, P Thibault, J Hébert, PP Roux, G Sauvageau

## Abstract

Given the particular attractivity of antibody-based immunotherapies, *in vitro* experimental approaches aiming to identify and quantify proteins directly located at the cell surface, such as the surfaceome, have been recently developed and improved. However, the “surface” enriched, yet noisy output obtained from available methods makes it challenging to accurately evaluate which proteins are more likely to be located at the surface of the plasma membrane and which are simple contaminants. To that purpose, we developed the *in silico* Surface Protein Annotation Tool (SPAT), which unifies established annotations to grade proteins according to the chance they have to be located at the cell surface. SPAT accuracy was tested using in-house acute myeloid leukemia data, as well as public datasets, and despite using publicly available annotations, showed good performances when compared to more complex surfaceome predictors. Given its simple input requirement, SPAT is easily usable for the annotation of any gene/protein lists. Its output, in addition to the “surface” score, provides additional annotations including a “secretion” flag, references to verified antibodies targeting annotated proteins, as well as expression data and protein levels in essential human organs, making it a user-friendly tool for the community.

## Introduction

It is estimated that nearly 20% of human genes encode cell surface proteins^1^ whose importance in cell biology, human disease, immunophenotyping, cell sorting and pharmacological targeting is well established. Unfortunately, a clear understanding of the full complement of cell surface proteins remains elusive because proteomic-based identification methods suffer from the hydrophobic nature of membrane proteins, which makes their purification difficult and subsequent detection problematic^2^. Recently developed chemoproteomic methods that target the cell-surface proteome (surfaceome) partly overcome this last issue^3,4^. However, the “surface” enriched, yet noisy, output obtained from these experiments makes it challenging to accurately discriminate surface proteins from other membrane-associated ones (e.g. components of the Golgi, endoplasmic reticulum, lysosome).

As for the *in silico* discrimination of cell surface from non-cell-surface proteins, predictors trained using different structural or biochemical features to identify such proteins have been developed^5,6^, however if they offer the possibility of *de novo* annotations, their classification relies on *in-house* models and do not necessarily match the subcellular localization already reported in curated databases. On the other hand, these databases represent multiple sources of information lacking standardization – both for localization information and confidence – making the automation of high-throughput data analysis particularly challenging and time-consuming. To facilitate this annotation of putative cell surface proteins from large-scale proteomic projects, we developed SPAT, a surface protein annotation tool that attributes a cell-surface score (from 0 to 10) by unifying previously established annotations from multiple curated proteomic databases.

In this manuscript, we evaluate the performance of SPAT using public datasets, surfaceome and total proteome datasets recently generated from acute myeloid leukemia (AML) cell lines, as well as transcriptome datasets from primary human AML specimens. We compare SPAT to other benchmark tools and demonstrate its potential to accurately annotate surface proteins.

## Material and Methods

### Implementation

SPAT scoring ranges from 0 to 10 according to the chance of a protein to be located at the cell surface (**Figure 1**). This score is calculated using annotations extracted from multiple databases including NeXtProt (https://www.nextprot.org/), GO (http://geneontology.org/), UniProt (https://www.uniprot.org/) and Human Protein Atlas (HPA, https://www.proteinatlas.org/). These annotations are classified in 6 “surface groups” (S1 to S5 and non-surface), S1 being the “surface” annotation group of highest confidence (**Sup. Table 1**). The algorithm then calculates an initial score based on the combinations of annotations/terms distributed into these groups. Finally, this initial score is weighted according to the eventual accumulation of “red” (associated with other cellular locations) or “secretion” (associated with extracellular release) annotation flags, lowering the chance of a protein to be found at the surface and hence its score (**Sup. Table 1**). As SPAT outputs the number of selected terms we associate with protein “secretion” (**Sup. Table 1**), for the sake of analyses, the count for each protein (scaled from 0 to 10) has been considered as a “secretion score”. As a sanity check, we tested the relation between SPAT score and basic topological information. As expected, high SPAT scores were enriched in proteins presenting several TM domains (**Sup. Figure 1A**), while high “secretion scores” were enriched in presence of a signal peptide (**Sup. Figure 1B**).

**Figure 1.**
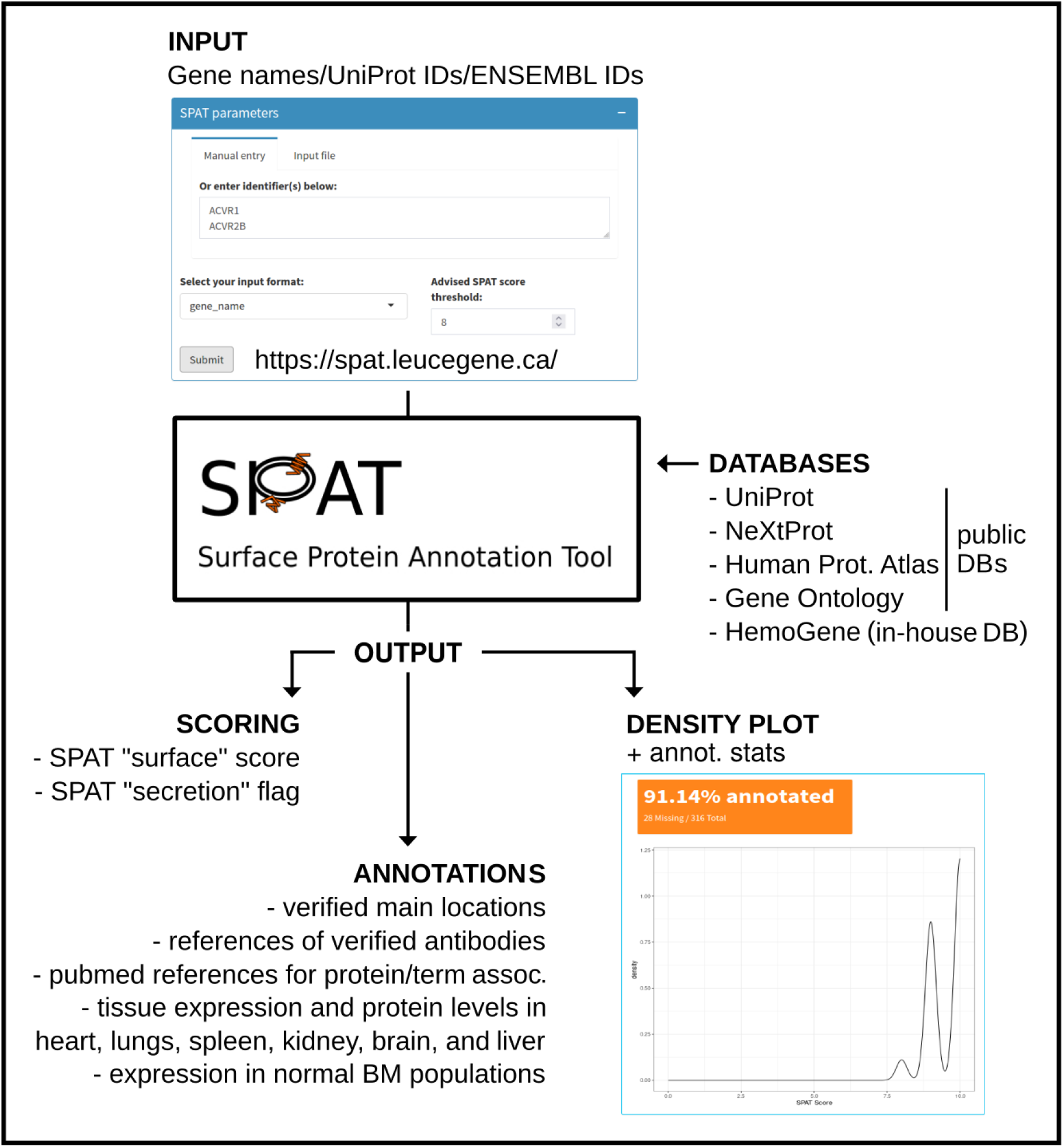
SPAT workflow.

In addition to the SPAT score and flags, the tool outputs a series of annotations from HPA and NeXtProt databases, including verified main locations (NeXtProt and HPA), references of verified antibodies (HPA), pubmed references for each protein/term association (NeXtprot), tissue expression and protein levels in heart, lungs, spleen, kidney, brain, and liver (HPA), as well as significant prognostic values for various tumors (HPA). Finally, a SPAT score distribution graph is available as part of the output.

### Cell culture

Human HL-60 (ATCC, CCL-240), KG1a (ATCC, CCL-246.1) and K562 (ATCC, CCL-3343) cell lines were cultured in DMEM supplemented with 10% FBS at 37°C in 5% CO2 and normal O2. Human NB4 cells (DSMZ, ACC 207) were cultured in RPMI 1640 supplemented with 10% FBS at 37°C in 5% CO2 and normal O2. Human OCI-AML3 and OCI-AML5 cell lines were obtained from Dr. Mark Minden at University Health Network, Toronto, Ontario, Canada through an MTA and were cultured in αMEM supplemented with 20% FBS with 10 ng/mL GM-CSF for OCI-AML5 at 37°C in 5% CO2 and normal O2. All cell lines were routinely tested using a mycoplasma contamination kit (R&D Systems).

### TMT labelling and LC-FAIMS-MS analysis

TMT labeling and FAIMS analysis were based on previously published methods^7^. Briefly, cells were chilled on ice, washed twice with cold PBS (pH 7.4), and pelleted by centrifugation (1000 rpm, 5 min). The pellets were resuspended in 1 mL of lysis buffer [8 M Urea, 50 mM HEPES pH 8.5, 75 mM sodium chloride, 1X protease inhibitor (complete, without EDTA, Roche), 1 mM sodium orthovanadate (Na3VO4), and 1 mM phenylmethylsulfonyl fluoride (PMSF)]. Cells were mechanically lysed by sonication. Cell debris and nuclei were removed by centrifugation at 17,000 x g for 5 min and proteins were reduced with 5 mM of TCEP for 30 min at room temperature then alkylated with 10 mM of 2-chloroacetamide at 37°C in the dark. 250 μg of lysate were precipitated using methanol/chloroform (lysate/methanol/chloroform/water 1:4:1:3 volume parts). Samples were vortexed and centrifuged for 5 min at 14,000 x g. The precipitated protein layer was washed twice with four volume parts of methanol and air-dried. For tryptic digestion, proteins were dissolved in 200 mM HEPES, pH 8.2 and trypsinized (enzyme/protein ratio 1:50) overnight at 37°C. Equal amounts of peptide aliquots were labeled with individual isobaric TMT10plex tags. Briefly, 100 μL of peptides (100 μg) dissolved in 200mM HEPES pH 8.2 were combined with 40 μL of TMT label (200 μg) in anhydrous acetonitrile for 90 min at room temperature. The reaction mixture was quenched with 1 μL of 50% hydroxylamine. The TMT tagged samples were equally mixed and then desalted using an Oasis HLB cartridge. The analysis by liquid chromatography-mass spectrometry, coupled to FAIMS (LS-FAIMS-MS) and data analysis were performed as described^1^.

### Cell surface protein enrichment and surfaceome analysis

The procedure for cell-surface protein enrichment was based on previously published methods^8^. Briefly, cells were chilled on ice, washed twice with ice-cold biotinylation buffer (PBS, pH 7.4, supplemented with 1 mM CaCl2 and 0.5 mM MgCl2), and incubated with 1 mg/mL Sulfo-NHS-LC-biotin (resuspended in biotinylation buffer) for 1 h at 4°C. The biotinylation reaction was quenched by addition of 100 mM glycine for 10 min at 4°C, followed by two washes with ice-cold biotinylation buffer. Biotinylated cells were incubated in surfaceome lysis buffer (SLB: 1% Triton X-100, 150 mM NaCl, 10 mM Tris–HCl, pH 7.6, 5 mM iodoacetamide (Sigma-Aldrich), 1× protease inhibitor (complete, without EDTA, Roche), 1 mM sodium orthovanadate (Na3VO4), and 1 mM phenylmethylsulfonyl fluoride (PMSF)) for 30 min at 4°C. Cell debris and nuclei were removed by successive centrifugation for 10 min at 4°C, initially at 2800 × g and then 16,000 × g. Next, biotinylated proteins were isolated from 1 mg total protein by incubating cell lysates with high-capacity streptavidin agarose resin (Thermo Fisher Scientific) for 2 h at 4°C. Beads were washed extensively with intermittent centrifugation at 1000 × g for 5 min to eliminate all potential contaminants bound to biotinylated proteins. Three washes were performed with SLB, once with PBS pH 7.4/0.5% (w/v) sodium dodecyl sulfate (SDS), and then beads were incubated with PBS/0.5% SDS/100 mM dithiothreitol (DTT), for 20 min at RT. Further washes were performed with 6 M urea in 100 mM Tris–HCl pH 8.5, followed by incubation with 6 M urea/100 mM Tris–HCl pH 8.5/50 mM iodoacetamide, for 20 min at RT. Additional washes were performed with 6 M urea/100 mM Tris–HCl pH 8.5, PBS pH 7.4 and then water. For proteomic analysis, beads were rinsed twice with 50 mM ammonium bicarbonate (NH4HCO3) pH 8.5, and re-suspended in 400 μL of 50 mM NH4HCO3 pH 8.5 containing 4 μg of proteomics grade trypsin (Sigma-Aldrich), overnight at 37°C. The proteins were further digested with an additional 4 μg trypsin for 4 h at 37°C. The resulting tryptic peptides were then collected by centrifugation at 10,000 × g, for 10 min at RT and the beads were washed twice with MS grade water. The tryptic fractions were dried to completion in a SpeedVac. Samples were then analyzed by mass spectrometry as described in Aubert *et al*., 2020^8^. Peptides were identified using PEAKS 7.0 (Bioinformatics Solutions, Waterloo, ON) and peptide sequences were blasted against the Human Uniprot database as previously described^8^.

### Membrane proteome “prediction datasets”

Membrane proteome datasets identified with different prediction methods (MEMSAT3^9^, MEMSAT-SVM^10^, Phobius^11^, SCAMPI^12^, SPOCTOPUS^13^, THUMBUP^14^, TMHMM^15^, and GPCRHMM^16^) were downloaded from the “Prediction of transmembrane protein topology and signal peptides” section (https://www.proteinatlas.org/humanproteome/tissue/secretome) of The human Protein Atlas database^17^.

### a-helical transmembrane domains and signal peptides information

The number of a-helical transmembrane (TM) domains and the presence of a signal peptide for each protein was obtained from The *in silico* human surfaceome dataset (http://wlab.ethz.ch/surfaceome/)^5^.

### Primary AML specimens

The Leucegene project is an initiative approved by the Research Ethics Boards of Université de Montréal and Maisonneuve-Rosemont Hospital. All leukemia samples were collected and characterized by the Quebec Leukemia Cell Bank after obtaining an institutional Research Ethics Board–approved protocol with informed consent according to the Declaration of Helsinki. The Quebec Leukemia Cell Bank is a biobank certified by the Canadian Tissue Repository Network. Detailed information on the cohort was previously published (see https://leucegene.ca/ and Moison *et al*. 2019^18^).

### HemoGene^19^ and CD34+CD45RA-cells expression data

Datasets are available in geo repositories: GSE98310 and GSE48846 (see “Data availability” section for detailed description of the samples and sequencing data location).

### Expression data processing

RNA-seq libraries were constructed according to TruSeq Protocols (Illumina) and sequencing was performed using an Illumina HiSeq 2000/4000 instrument. Trimming of sequencing adapters and low-quality bases was done using the Trimmomatic (v0.38) tool^20^. Resulting reads were mapped to the reference using the RNA-seq aligner STAR (v2.7.1)^21^ and quantification of gene and isoform expression were performed using RSEM (v1.3.2)^22^. Statistical analyses of all experiments were done using R. The limma package (v3.28.14) and its Voom method^23^ were used to conduct differential expression analysis and obtain gene significance.

### GO term enrichments

GO (http://geneontology.org/) significant enrichments were conducted using the GOrilla tool^24^. Target sets were composed of genes coding for proteins associated with a SPAT score of interest, while background sets were composed of remaining coding genes.

## Results

### SPAT distinguishes surface *versus* intracellular membrane proteins

The accurate *in silico* annotation of the subcellular localization of proteins remains a challenge. Numerous predictive methods are dependent on training with “peptide” features such as amino acid physicochemical properties. These usually efficiently identify membrane proteins, but often fail to accurately discriminate cell surface proteins from proteins located at the membrane of intracellular organelles such as the Golgi apparatus or the endoplasmic reticulum (ER).

To evaluate the ability of SPAT to distinguish intracellular membrane proteins from cell surface proteins, we graded membrane proteins obtained from “prediction datasets” generated by 7 human membrane proteome prediction tools that do not specifically aim to distinguish cell surface from intracellular proteins (MEMSAT3, MEMSAT-SVM, Phobius, SCAMPI, SPOCTOPUS, THUMBUP and TMHMM) and, for each of these algorithms, we plotted the proportion of predicted proteins at each SPAT score unit (**Figure 2A, Supplemental methods**).

**Figure 2.**
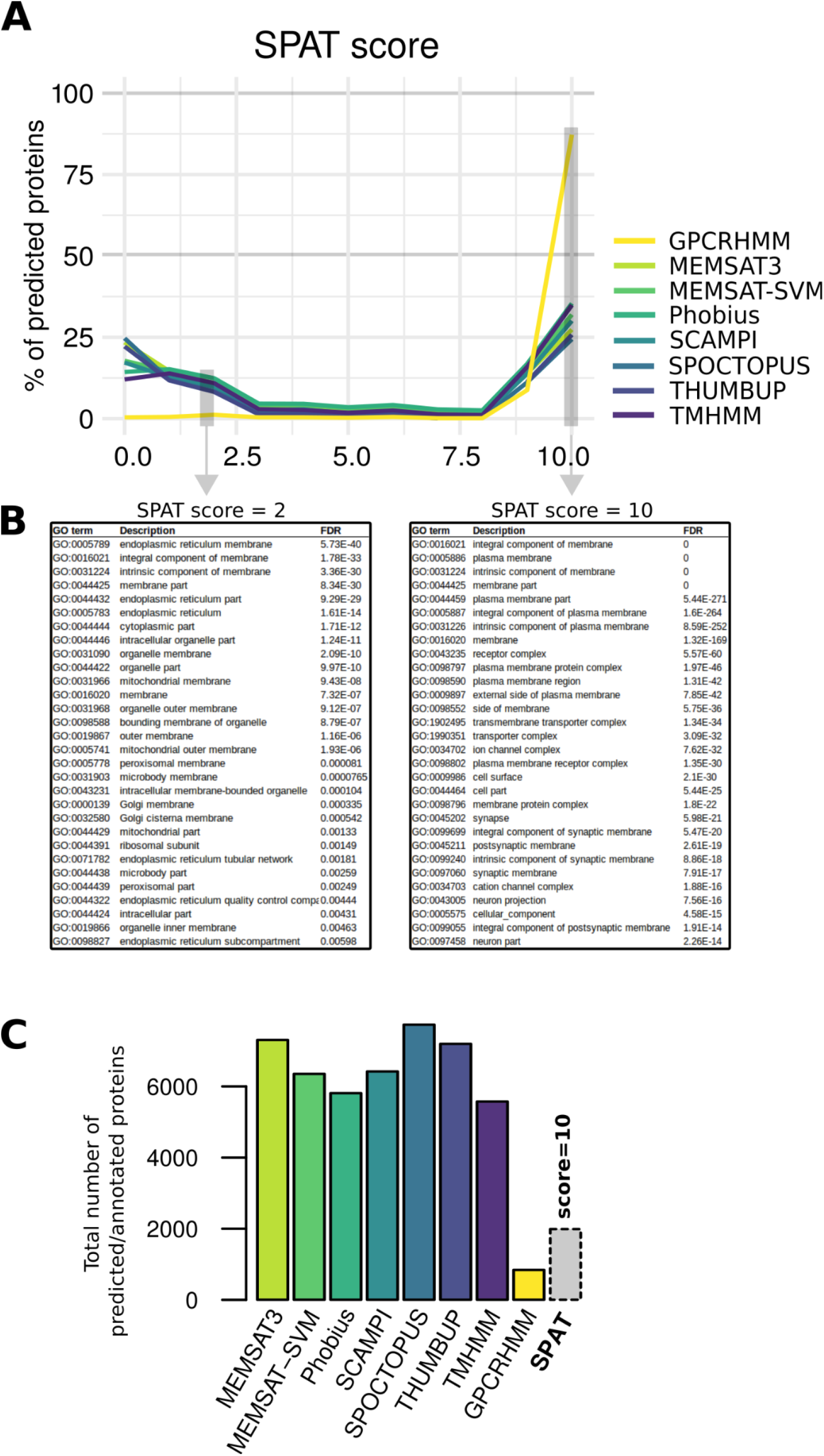
**A**. SPAT score distribution of predicted membrane proteins by each tested algorithm. At each SPAT score unit is plotted the proportion (%) of predicted proteins by each algorithm (as indicated in the legend, each curve represents data relative to one algorithm). **B**. GO term enrichments calculated using all predicted proteins associated with a SPAT score of 2 or 10 (left and right table, respectively) as input. **C**. Total number of predicted membrane proteins for each membrane proteome predictor tested. The color code matches the legend of panel A. As an indicative value, the total number of proteins associated with a SPAT score of 10 is also represented (gray bar with a dotted border).

We also analyzed the output obtained from GPCRHMM, which uses a hidden Markov model that exploits the common topology of G protein-coupled receptors (GPCRs) to specifically predict cell surface receptors from amino acid sequence. GO enrichment analysis of predicted proteins presenting low and high SPAT scores (2 and 10) revealed strong enrichments of “intracellular membrane” and “cell surface proteins”-associated terms, respectively, demonstrating that SPAT efficiently discriminates between these two types of membrane proteins (**Figure 2B**). The strong enrichment of GPCRHMM-identified proteins observed at high SPAT scores (starting at score = 9) confirmed that SPAT is able to correctly grade GPCRs. While GPCRHMM limits its prediction to this class of surface proteins, which explains the very limited number of classified entities (n = 753, **Figure 2C**), SPAT grading is independent of protein type. As a result, with > 19,000 genes considered by SPAT analysis, SPAT annotated 2,002 genes coding for proteins with the highest confidence score of 10 (**Figure 2C**).

### Surfaceome data, unlike total proteome, show enrichment of proteins with high SPAT scores

To estimate the performance of our algorithm with experimental data, we conducted both surfaceome and global proteome analysis of a pool of 6 different cell lines (OCI-AML3, OCI-AML5, NB4, HL60, KG1a and K562) and scored the output of each experimental approach using SPAT, expecting an important enrichment of proteins with high SPAT scores in surfaceomes *versus* global proteomes (SPAT annotation outputs: **Sup. Table 2**). While scores obtained for the total proteome experiment showed an unimodal distribution with a peak at low SPAT scores (Mo #1 = 1, **Sup. Figure 2A**) indicative of non surface proteins, a bimodal distribution SPAT scores was found for the surfaceome dataset (**Sup. Figure 2B**) in which there was an important enrichment of proteins with high SPAT scores forming a second peak (Mo #2 = 9.5). SPAT score analysis thus expectedly suggests a strong enrichment of surface proteins in surfaceome *versus* total proteome datasets.

### Comprehensive comparison between SPAT and a benchmark surfaceome predictor

To evaluate the accuracy of our algorithm, we compared its performance to the machine-learning-based surfaceome predictor SURFY^5^, a reference tool specifically selected because of its high performance. SURFY’s predictions (graded from 0 to 1) are made by a random forest classifier trained using topological information of experimentally verified high-confidence cell-surface proteins. These predictions are restricted to ∼7,900 proteins (5,500 a-helical TM proteins + a selection of 2,403 proteins without the domain), resulting in the absence of grading for a series of entries, some of which reaching high SPAT scores (e.g. n = 128 ungraded proteins by SURFY with a SPAT score ⩾ 8). While the distribution of scores generated by both methods showed a common trend (**Sup. Figure 3A**, Pearson’s r = 0.69), especially at low and high values, the limited correlation reflected underlying differences in the “surface proteins” evaluation.

In order to conduct a fair and qualitative comparison of the two algorithms, and given differences in score origin and scale between the two methods, we tested different SPAT score thresholds against SURFY’s authors chosen threshold (⩾ 0.5818 for “a representative human surfaceome”^5^) using surfaceome and global proteome data presented in the previous section, and retained the SPAT score allowing for a similar enrichment of proteins located at the cell surface for the two methods (**Figure 3A and Sup. Figure 3B**). Interestingly, resulting SPAT threshold (⩾ 8) corresponds to the score we usually consider as optimal for surfaceome analysis, i.e. limiting both false positives and negatives, as the analysis of cluster of differentiation (CD) proteins presented below pointed it out. This threshold also corresponds to the first score unit located after the inflection point of the “surface” enrichment peak (#2) obtained using the surfaceome dataset (**Sup. Figure 2B**).

**Figure 3.**
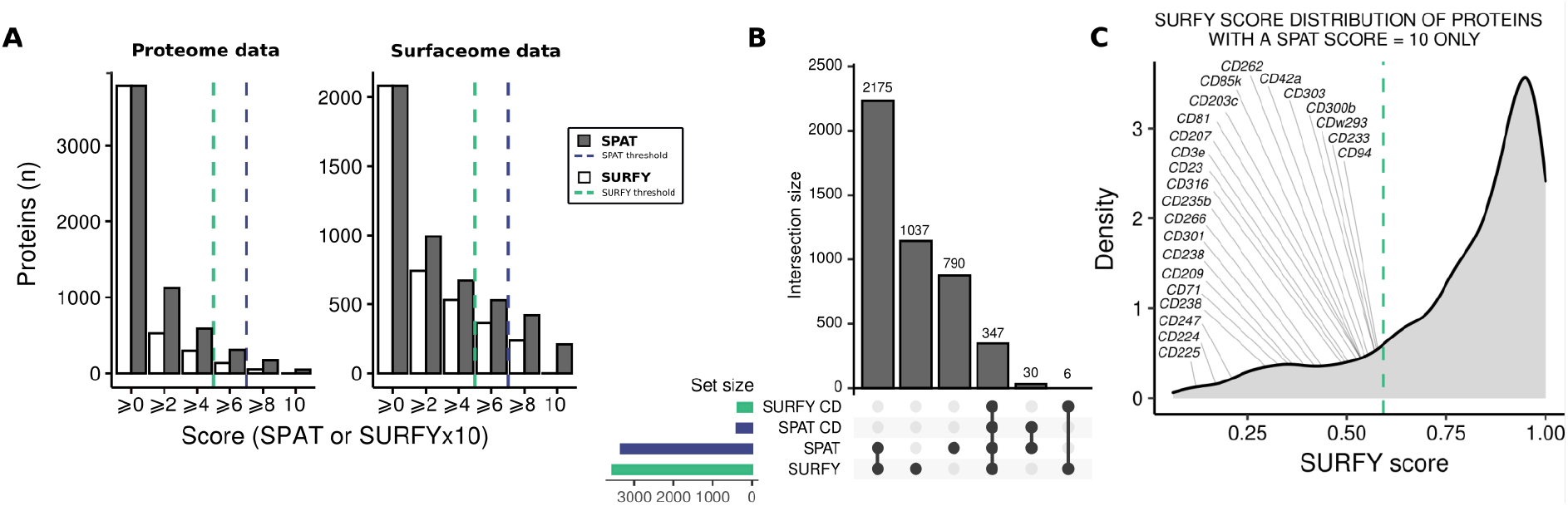
**A**. Distribution of SPAT (dark bars) and SURFY (white bars) scores calculated for global proteome (left panel) and surfaceome (right panel) data obtained from a mix of 6 different cell lines (OCI-AML3, OCI-AML5, NB4, HL60, KG1a and K562). SURFY and SPAT thresholds are indicated by green (⩾ 0.5818×10) and blue (⩾ 8) dotted lines, respectively. **B**. UpSet representation of “surface proteins” classification by SURFY (green bar) and SPAT (blue bar) using previously defined thresholds. Considered sets and intersections are indicated by dots and links between dots, respectively, at the bottom of the figure. “SPAT CD” and “SURFY CD” correspond to sets of Cluster of Differentiation proteins passing algorithms’ thresholds. **C**. SURFY score distribution of all proteins presenting a SPAT score = 10. Author’s chosen threshold for SURFY (⩾ 0.5818) is indicated by a green dotted line. SURFY scores obtained for CD proteins with a SPAT score = 10 are indicated directly on the curve.

Using these thresholds, SURFY and SPAT commonly classified close to 2,200 proteins as surface proteins, while 1,037 and 790 additional proteins were specific to each method, respectively (**Figure 3B**). In their article, SURFY’s authors considered CD proteins as *bona fide* cell surface proteins and used them as an indicator of performance for the classification. Here, 347 CD proteins passed both SURFY and SPAT confidence thresholds but interestingly, SPAT classified 30 additional CDs as “surface proteins” against 6 for SURFY (**Figure 3B and 3C, Sup. Table 3 and 4**). Of note, 23 of the “SPAT-specific” CDs received a maximum SPAT score of 10 when SURFY scores for these proteins ranged from 0.1517 (CD225) to 0.5808 (CD94) (**Figure 3C, Sup. Table 4**). And while all of the 6 “SURFY-specific” CD proteins presented either direct (detection) or indirect (prediction) evidence against a main location at the plasma membrane in databases such as UniProtKB/Swiss-Prot, COMPARTMENTS and/or HPA (**Sup. Table 4**), making them ambiguous “surface” candidates, the majority of the 30 “SPAT-specific” CDs (n = 20) presented annotations that suggested or even confirmed their location at the cell membrane (**Sup. Table 4**), justifying their classification as putative surface proteins by SPAT. For example, CD72, which was recently reported as a target for CAR-T cells in *KMT2A*/*MLL1*-rearranged B-ALL^25^, received a SPAT score of 9 whereas the protein was classified as non-surface by SURFY with a score of 0.29. Overall, these results further demonstrate the accuracy of SPAT annotations.

### Example of SPAT annotations of cell type-specific surface and secreted markers

To illustrate the interest of our method in a more specific example, we integrated SPAT annotations to an analysis aiming to identify cell type-specific surface and secreted markers from transcriptome data. We exploited the HemoGene dataset previously published by our group (**Supplemental methods**), which provides reference expression value for normal human haematopoietic cell subsets. Using two sets composed of the top 30 genes with the highest expression variance in the dataset and coding for proteins either annotated at the cell surface (SPAT score ⩾ 8, no “secretion” flag, **Figure 4A and 4B**), or presumably secreted (conservative SPAT “secretion” score threshold ⩾ 5, corresponding to the 98^th^ percentile of the “secretion” score distribution, **Figure 4C, 4D, and Sup. Figure 4**), we were able to obtain logical and comparable clustering of HemoGene samples (**Figure 4B and 4D**).

**Figure 4.**
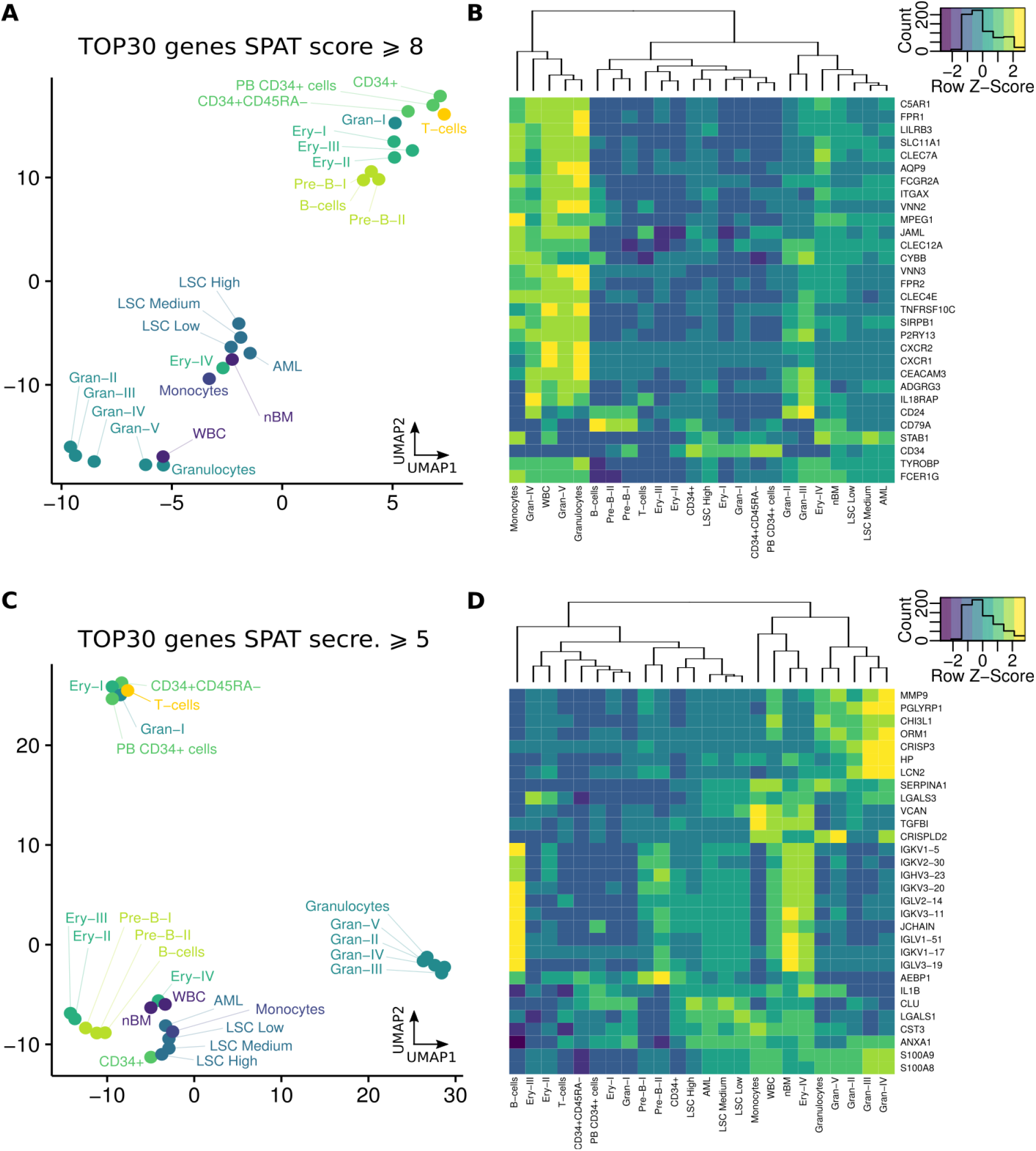
**A**. UMAP projection of HemoGene samples based on the expression (TPM) of the 30 most variable genes (based on variance) presenting a SPAT score ⩾ 8 (“secretion” score = 0). **B**. Expression heatmap for the corresponding “surface” genes (right). Samples (bottom) are ordered according to the dendrograms (top) obtained from an unsupervised hierarchical clustering. **C**. UMAP projection of HemoGene samples based on the expression (TPM) of the 30 most variable genes (based on variance) presenting a SPAT “secretion score” ⩾ 4. **D**. Expression heatmap for the corresponding “secretion” genes (right). Samples (bottom) are ordered according to the dendrograms (top) obtained from an unsupervised hierarchical clustering (see “Data availability” section for detailed description of the samples and sequencing data location).

SPAT-annotated “surface” dataset presented expected and widely used cell markers – such as the progenitor-associated CD34 transmembrane phosphoglycoprotein^26,27^, CD79A in B-cells^28^, ITGAX (CD11c) in monocytes and granulocytes^29^, CD24 in pre-B cells and granulocytes^30,31^ – but also more recently identified markers such as ADGRG3^32^, VNN2 (GPI-80)^33^ or MPEG1^34^. Of note, SPAT correctly identified the G-protein coupled receptor P2RY13, previously identified as overexpressed in AML^20^, as a potential granulocyte surface marker. Similarly to this *in silico* surfaceome, SPAT-defined secretome associated with HemoGene samples (**Figure 4D**) reported well-known factors, among which are the calcium binding factors S100A8 (MRP8) and S100A9 (MRP14), abundantly produced by myeloid cells^35^, and the neutrophil gelatinase-associated lipocalin LCN2 (NGAL)^36^.

### Example of SPAT annotations of antigens targeted by immunotherapies

The previous section demonstrated the potential of SPAT annotation for marker exploration using expression data and suggested that our algorithm could be a useful assistant for the identification of cancer cell markers and immunotherapeutic targets. To further investigate SPAT annotation performance on targetable surface antigens, we analyzed human targets registered in The Therapeutic Structural Antibody Database^37^ (Thera-SAbDab) (**Figure 5A and Sup. Table 5**) and compared it to SURFY’s analysis. Both tools proposed a rather accurate classification of most antigens with 201/296 commonly and rightly identified as surface proteins. Once again, expectedly given the SPAT scoring method, the majority of “SPAT-specific” surface classification (n = 18, **Figure 5A**) were justified by annotations available in several public databases (**Sup. Table 5**).

**Figure 5.**
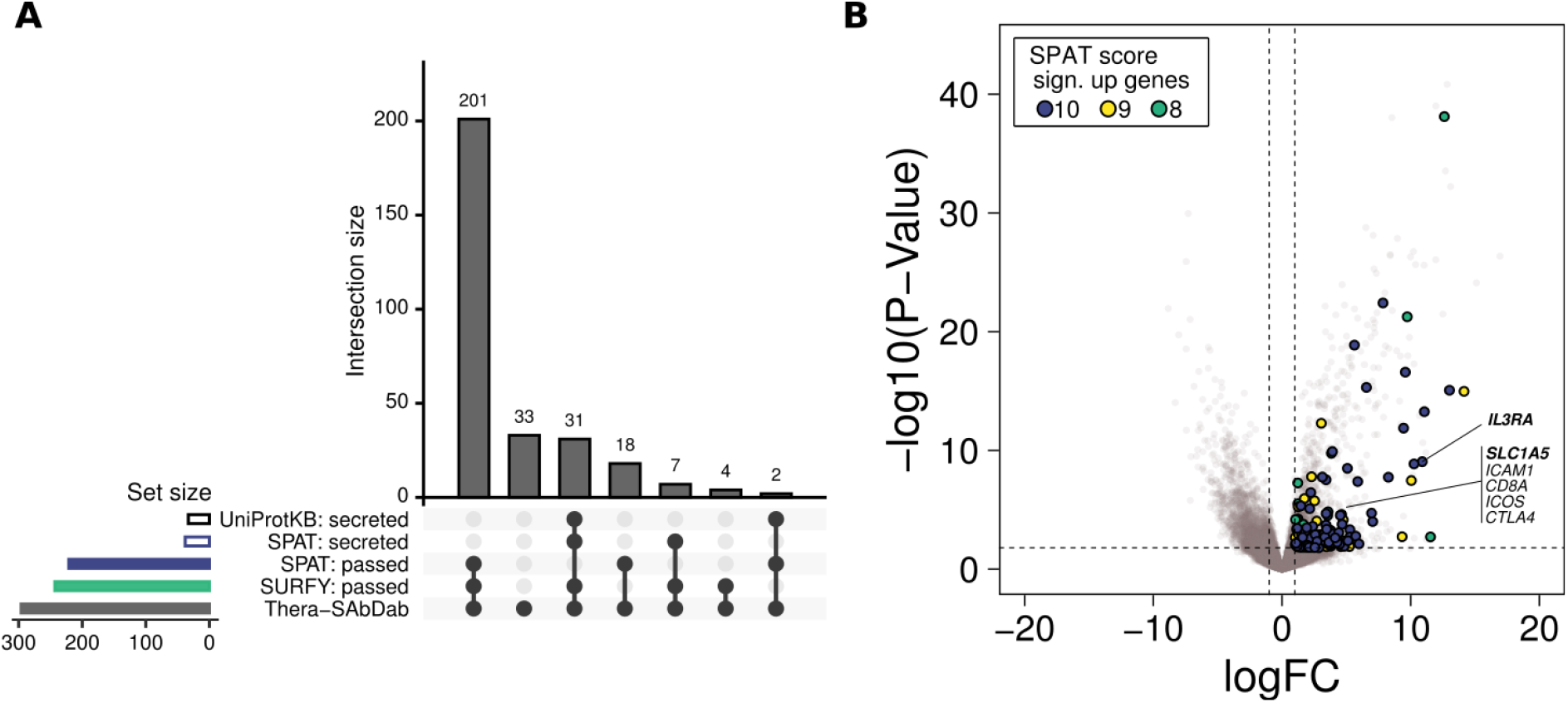
**A**. UpSet representation of SPAT and SURFY classification of human targets registered in The Therapeutic Structural Antibody Database (Thera-SAbDab, grey bar, n = 296). Considered sets and intersections are indicated by dots and links between dots, respectively, at the bottom of the figure. “SPAT: passed” (blue bar) and “SURFY: passed” (green bar) correspond to proteins classified as “surface” proteins by each algorithm using previously defined thresholds. “SPAT: secreted” proteins (empty blue bar) correspond to proteins with a SPAT score < 8 and presenting a SPAT “secretion” flag (SPAT “secretion score” ⩾ 1, Sup. Table 5). “UniProtKB: secreted” proteins (empty black bar) correspond to proteins for which “secreted” is the only location identified in the UniProtKB/Swiss-Prot database (Sup. Table 5). **B**. Volcano plot representation of the differential expression analysis conducted on RNAseq data comparing complex karyotype AML (n = 68, see “Data availability” section for detailed description of the samples and sequencing data location) *versus* CD34+CD45RA-cord blood cell specimens (n = 5, see “Data availability” section for detailed description of the samples and sequencing data location). The horizontal dashed line indicates an adjusted p-value of 0.05 and vertical dashed lines indicate log fold change (logFC) of 1 and -1. Green, yellow and blue dots correspond to genes significantly overexpressed (logFC > 1, FDR < 0.05) and presenting a SPAT score = 8, = 9 and = 10, respectively. Annotated genes code for cancer target proteins referenced in Thera-SAbDab (in bold for AML targets).

SPAT also properly classified 31 antigens as putatively secreted – using the presence of a unique “secreted” location in UniProtKB/Swiss-Prot as reference annotation (**Sup. Table 5**) – while, on the other hand and as suspected by SURFY’s authors, their model failed to accurately distinguish secreted from surface proteins in absence of a-helical TM domains^5^, which was the case for all of these 31 proteins (**Sup. Table 5**).

To further assess SPAT annotation of targetable antigens, we simulated a “real data” example by conducting a differential expression analysis between dismal prognosis complex karyotype (CK) AML specimens (defined as presenting 3 and more chromosomal abnormalities) from the Leucegene cohort and CD34+CD45RA-cord blood cells obtained from 5 non-pooled individuals (**Supplemental methods, Figure 5B, Sup. Table 6**). SPAT annotations of significantly overexpressed genes in CK AML specimens identified IL3RA (CD123), already studied as an immunotherapy target for refractory AML, as well as several surface proteins targeted by antibodies referenced in Thera-SAbDab which could represent as many valuable targets for the development of immunotherapy against AML cells.

Overall, these results further confirmed SPAT as an asset for easy data annotation facilitating the exploitation of diverse analysis outputs.

## Discussion

Cell surface proteins represent attractive drug targets. Recently, immunotherapeutic approaches exploiting the overexpression of specific cell surface antigens by malignant cells highlighted the need to easily and accurately annotate the surfaceome. Existing tools such as SURFY^5^ and others^6^ use protein domain information and associated biochemical features in machine learning approaches to predict subcellular localization of membrane proteins.

While these methods offer the advantage of *de novo* predictions of surfaceome proteins – which is a limitation of our tool given that it relies on pre-existing annotations – results remain highly dependent on training limitations (e.g. representativity, quality of datasets) and can present a lack of accuracy when compared to curated information. Furthermore, the annotation is often limited to a subset of pre-selected proteins, which is somewhat opposed to the exploratory interest of such approaches. Overall, these approaches, although innovative and at least partly independent of existing annotations, may not be appropriate for projects needing a simple evaluation of protein localization based on up to date and curated information.

In this case, SPAT can offer a user-friendly and reliable alternative by unifying the information obtained from multiple databases (using curated and regularly updated selection of annotations) and facilitating its interpretation by retrieving a simple representative score of the likelihood a protein has to be present on the cell surface. Note that the purpose of SPAT annotation is not to take into account specific, transient and transformed cellular states which can lead to non-canonical localizations, but rather to provide annotations depicting a general biological context.

The simplified web interface allows users to easily identify and annotate surface proteins from any kind of data by simply uploading a list of proteins/genes in several formats including UniProt and Ensembl IDs. Indeed, as SPAT only requires a list of gene IDs as input, this tool can easily be integrated in projects exploiting other kinds of omics, such as transcriptomic data.

Furthermore, SPAT proposes a series of additional annotations including, among others, references to verified antibodies targeting annotated proteins as well as expression data and protein levels in essential human organs, making it a useful tool for the identification of immunotherapy targets. This is especially true given that, as opposed to numerous other methods, SPAT annotations are available for more than 19,000 protein coding genes and not limited to a specific subclass of proteins.

## Author Contributions

J.F.S. contributed to project conception, wrote the algorithm, analyzed the data, generated all figures and wrote the manuscript, L.T. contributed to project conception, surfaceome data generation and algorithm design, L.A. contributed to project conception and surfaceome data generation, E.A. and G.B. contributed to the interface conception and maintenance, S.P. contributed to proteome data generation, E.B. contributed to surfaceome and proteome data generation, M.E.B. contributed to project conception, P.T. supervised proteome data generation, J.H. contributed to project conception, provided AML samples and clinical data of the Leucegene cohort, P.P.R. contributed to project conception and supervised surfaceome data generation, G.S. contributed to project conception, coordination and supervision.

## Acknowledgements

The authors wish to thank Muriel Draoui for project coordination. J.F.S. was supported by an IVADO and Canada First Research Excellence Fund (Apogée/CFREF) and a Canadian Institutes of Health Research (CIHR) postdoctoral fellowships. This work was supported by the Government of Canada through Genome Canada and the Ministère de l’économie et de l’innovation du Québec through Génome Québec (ref. grant #4524 and grant #13528). J.H. holds a research chair from Industrielle-Alliance (Université de Montréal). BCLQ is supported by grants from the Cancer Research Network of the Fonds de recherche du Québec–Santé. RNA-Seq read mapping and transcript quantification were performed on the supercomputer Briaree from Université de Montréal, managed by Calcul Québec and Compute Canada. The operation of this supercomputer is funded by the Canada Foundation for Innovation (CFI), NanoQuébec, RMGA and the Fonds de recherche du Québec - Nature et technologies (FRQ-NT).

## Data availability

SPAT algorithm is available at https://github.com/sauvageau-lab/SPAT. The user interface is available at spat.leucegene.ca.

Detailed description of the samples and sequencing datasets are available in GEO repositories listed below.

- HemoGene data: https://www.ncbi.nlm.nih.gov/geo/query/acc.cgi?acc=GSE98310

- CD34+CD45RA-cord blood cell specimens (ED11, ED14, ED15, ED18 and ED31): https://www.ncbi.nlm.nih.gov/geo/query/acc.cgi?acc=GSE48846

- AML specimens: https://www.ncbi.nlm.nih.gov/geo/query/acc.cgi?acc=GSE67040

## Supplemental data

This article contains supplemental data.

