## Supplemental data for "SPAT: Surface Protein Annotation Tool"

Affiliations: 1. Institute for Research in Immunology and Cancer, Université de Montréal, Montréal, Québec, Canada. 2. Department of Chemistry, Université de Montréal, Montréal, Canada. 3. Leukemia Cell Bank of Quebec, Maisonneuve-Rosemont Hospital, Montréal, Canada. 4. Division of Hematology, Maisonneuve-Rosemont Hospital, Montréal, Québec, Canada. 5. Department of Medicine, Faculty of Medicine, Université de Montréal, Montréal, Canada. 6. Department of Pathology and Cell Biology, Faculty of Medicine, Université de Montréal, Montréal, Canada.

### Supplemental Figure legends

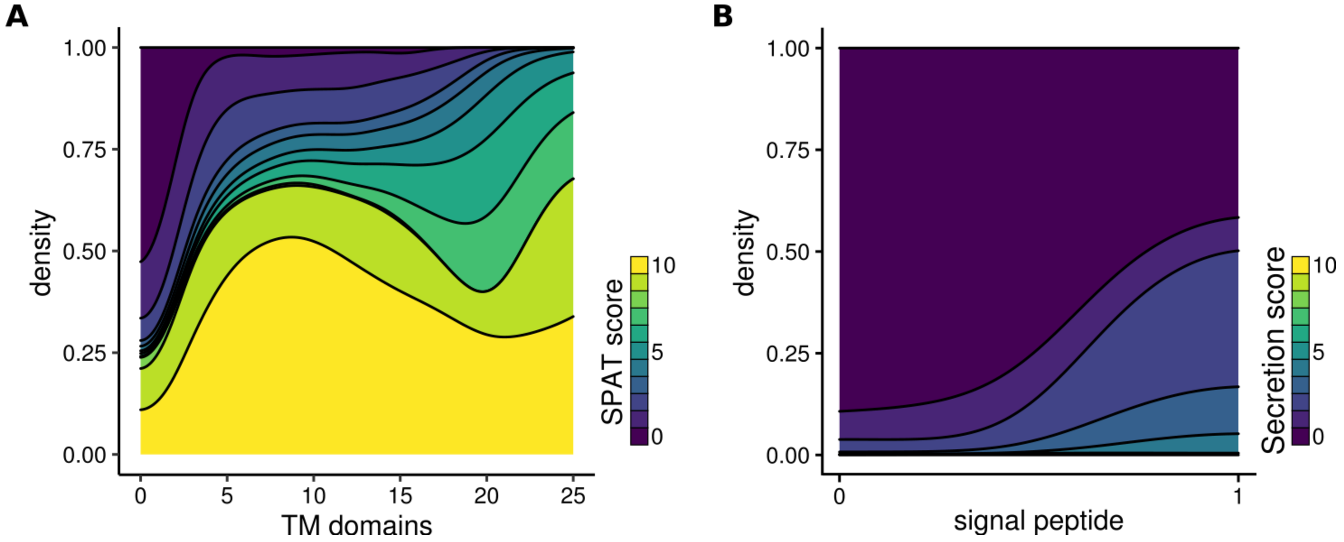


**Sup. Figure 1. A.** SPAT score distribution (stacked density) according to the number of α-helical transmembrane (TM) domains per protein. **B.** SPAT "secretion score" distribution (stacked density) according to the presence of a signal peptide.

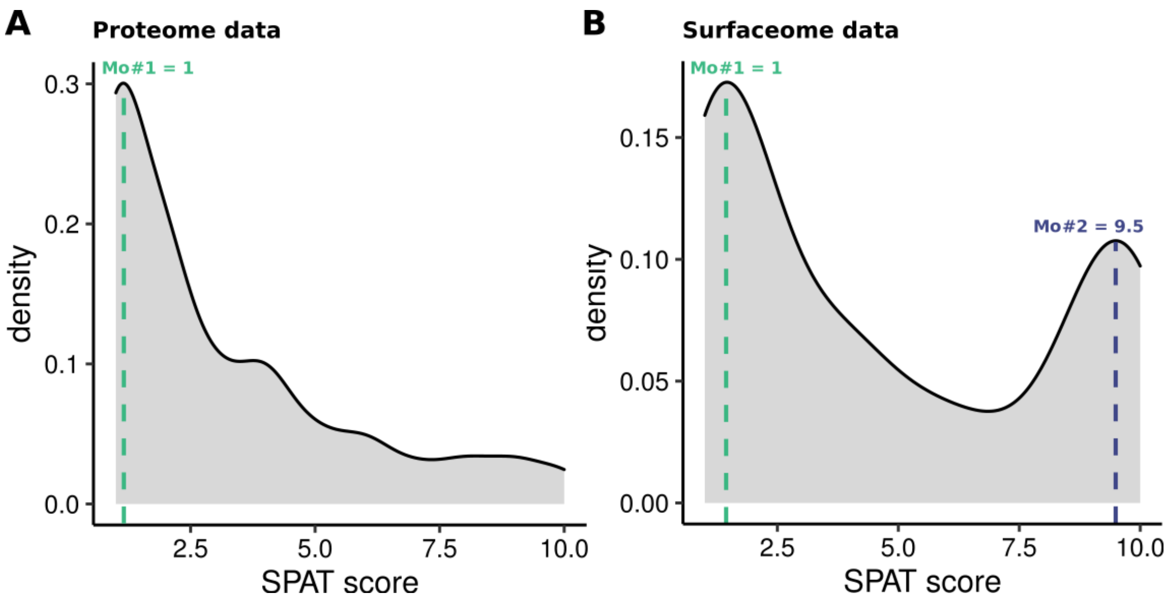


**Sup. Figure 2.** Distribution of SPAT scores calculated for global proteome (**A**) and surfaceome (**B**) data obtained from a mix of 6 different cell lines (OCI-AML3, OCI-AML5, NB4, HL60, KG1a and K562). The modes of distributions (Mo #1 and Mo #2) are indicated by green and blue dotted lines, respectively.

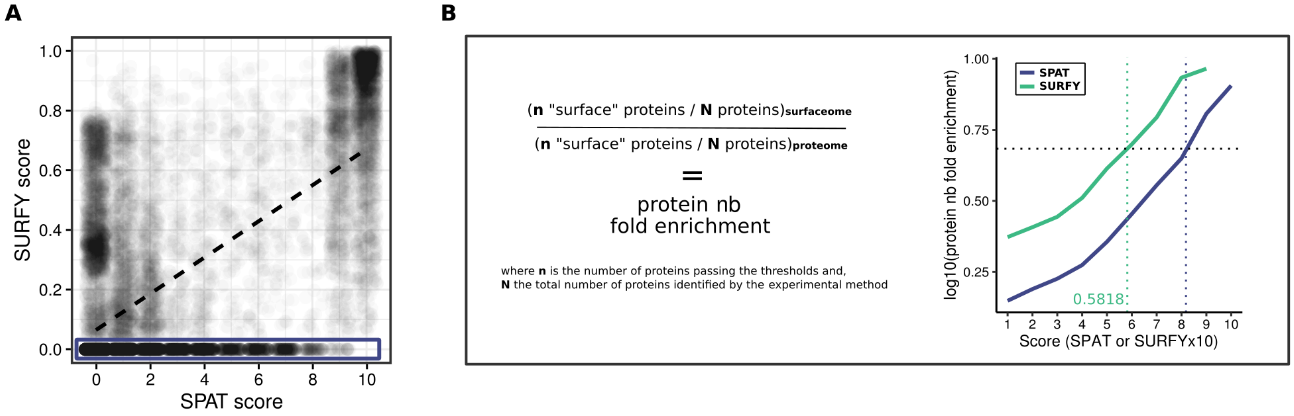


**Sup. Figure 3. A.** Comparison of SPAT *vs.* SURFY scores. The dotted black line corresponds to the linear regression line. The blue frame indicates proteins graded by SPAT but ungraded by SURFY. **B.** Calculation of the protein number fold-changes for different SPAT and SURFY score thresholds in a surfaceome *vs.* proteome comparison (**left panel**) and corresponding enrichment curves calculated for SPAT (solid blue line) and SURFY (solid green line) (**right panel**). Vertical dotted green and blue lines indicate SURFY's chosen threshold and corresponding SPAT threshold, respectively, to obtain a comparable enrichment value depicted by an horizontal black dotted line.

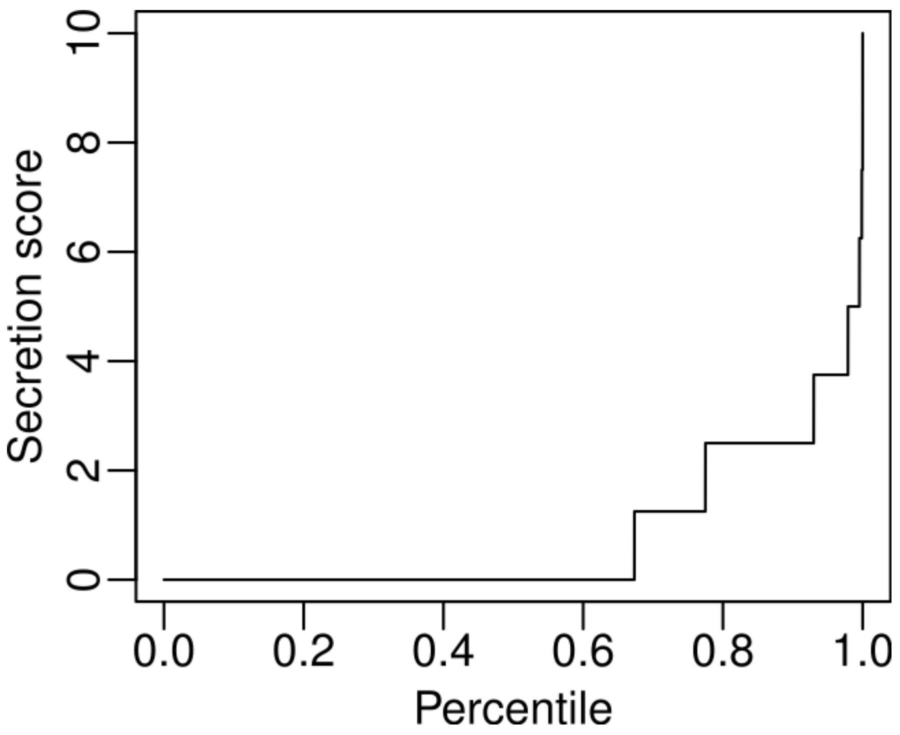


**Sup. Figure 4.** Cumulative percentiles of annotated proteins according to SPAT “secretion” score.

### Supplemental Tables

**Sup. Table 1.** Annotation terms and classification.

**Sup. Table 2.** SPAT annotation of global proteome and surfaceome data obtained from a mix of 6 different cell lines (OCI-AML3, OCI-AML5, NB4, HL60, KG1a and K562), as produced by the interface.

**Sup. Table 3.** SPAT annotation of CD proteins, as produced by the interface.

**Sup. Table 4.** SPAT and SURFY-specific CD proteins.
For annotations extracted from COMPARTMENTS and The Human Protein Atlas, information into brackets indicate the level of reliability of the annotation, see respective database websites for details.

**Sup. Table 5.** SPAT and SURFY-specific Thera-SAbDab targets.
For annotations extracted from COMPARTMENTS and The Human Protein Atlas, information into brackets indicate the level of reliability of the annotation, see respective database websites for details.

**Sup. Table 6.** Output of differential expression analysis conducted on RNAseq data comparing complex karyotype AML specimens from the Leucegene cohort (n=72) and CD34+CD45RA- cord blood cell samples obtained from non-pooled individuals (n=5).
